# Sleep loss disrupts the neural signature of successful learning

**DOI:** 10.1101/2021.11.16.468870

**Authors:** Anna á V. Guttesen, M. Gareth Gaskell, Emily V. Madden, Gabrielle Appleby, Zachariah R. Cross, Scott A. Cairney

## Abstract

Sleep supports memory consolidation as well as next-day learning. The influential *Active Systems* account of offline consolidation suggests that sleep-associated memory processing paves the way for new learning, but empirical evidence in support of this idea is scarce. Using a within-subjects (N = 30), crossover design, we assessed behavioural and electrophysiological indices of episodic encoding after a night of sleep or total sleep deprivation in healthy adults (aged 18-25 years), and investigated whether behavioural performance was predicted by the overnight consolidation of episodic associations formed the previous day. Sleep supported memory consolidation and next-day learning, as compared to sleep deprivation. However, the magnitude of this sleep-associated consolidation benefit did not significantly predict the ability to form novel memories after sleep. Interestingly, sleep deprivation prompted a qualitative change in the neural signature of encoding: whereas 12-20 Hz beta desynchronization – an established marker of successful encoding – was observed after sleep, sleep deprivation disrupted beta desynchrony during successful learning. Taken together, these findings suggest that effective learning depends on sleep, but not necessarily sleep-associated consolidation.

## Introduction

How do we remember events from days gone by? It is now firmly established that sleep facilitates memory consolidation; the process by which weak and initially labile memory traces become strong and enduring representations (Ashton and Cairney 2021; Ashton et al. 2020; Cairney et al. 2018b; Durrant et al. 2016; Gais et al. 2006; Gaskell et al. 2018; Payne et al. 2012; Talamini et al. 2008). Whereas sleep was originally thought to provide only passive protection to memory consolidation (i.e. by shielding memories from the interference posed by wakeful experience), recent work suggests that newly formed memories are actively strengthened during sleep (Cairney et al. 2018a; Rasch et al. 2007; Schönauer et al. 2017; Schreiner et al. 2021; Wang et al. 2019).

The influential *Active Systems* account of sleep-associated consolidation posits that the reactivation of hippocampus-dependent memories during slow-wave sleep (SWS) facilitates their migration to neocortex for long-term storage (Born and Wilhelm 2012; Klinzing et al. 2019; Rasch and Born 2013; Walker 2009). Supporting this view, functional neuroimaging studies have shown that overnight consolidation supports a shift in the memory retrieval network from hippocampus to neocortex (Takashima et al. 2009), with time spent in SWS predicting the reduction in hippocampal retrieval dependency (Cairney et al. 2015; Takashima et al. 2006). Along the same lines, other work has shown that post-learning sleep (as compared to sleep deprivation) promotes functional coupling between activity in the hippocampus and prefrontal cortex when retrieval is assessed 48 h later (Gais et al. 2007). Taken together, these findings suggest that hippocampal-to-neocortical information transfer emerges during the first nights after learning, although the consolidation process presumably takes many weeks or even months to complete (Dudai 2004; Dudai et al. 2015).

While the benefits of sleep for memory consolidation are well known, recent work has indicated that sleep also supports next-day learning of hippocampus-dependent memories. When a night of sleep deprivation precedes a novel learning opportunity, declarative memory recall is severely impaired, even after recovery sleep (Alberca-Reina et al. 2014; Cousins et al. 2018; Kaida et al. 2015; Tempesta et al. 2016), suggesting that an absence of sleep disrupts memory encoding in hippocampus. Indeed, as compared to a normal night of sleep, sleep deprivation weakens hippocampal responses during successful learning (i.e. for memories that are correctly recalled in a later retrieval test, after recovery sleep), leading to an overall decline in recall performance (Yoo et al. 2007). Correspondingly, daytime naps not only facilitate learning (Mander et al. 2011), but also restore hippocampal encoding capabilities, as compared to an equivalent period of wakefulness (Ong et al. 2020).

The interplay of various brain rhythms has been identified as a key mechanism that regulates communication between hippocampus and neocortex during sleep-associated memory processing. Slow oscillations (< 1 Hz EEG activity) have been causally linked to overnight memory retention (Leminen et al. 2017; Marshall et al. 2006; Ngo et al. 2013; Ong et al. 2016; Papalambros et al. 2017; Perl et al. 2016) and are thought to play a central role in the reactivation and reorganisation of hippocampus-dependent memories (Born and Wilhelm 2012; Klinzing et al. 2019; Rasch and Born 2013; Walker 2009). Delta waves (1-4 Hz), by contrast, have been implicated in forgetting via processes of synaptic renormalization (Genzel et al. 2014) and are thought to interact with slow oscillations to regulate the balance between memory consolidation and weakening (Kim et al. 2019).

Intriguingly, neural oscillations implicated in overnight memory processing have also been linked to new learning in hippocampus, suggesting that these processes rely on overlapping mechanisms. For example, selectively suppressing slow-wave activity SWA (0.5-4 Hz) via an acoustic perturbation approach impairs declarative memory encoding and reduces encoding-related activity in hippocampus (Van Der Werf et al. 2009). Reciprocally, enhancing SWA though electrical stimulation improves encoding of hippocampus-dependent memories but not non-hippocampal procedural skills (Antonenko et al. 2013). Augmenting slow oscillations via auditory stimulation leads to similar effects, with the magnitude of the slow oscillation enhancement predicting both hippocampal activation and behavioural performance at encoding (Ong et al. 2018). To what extent memory processes mediated by sleeping brain rhythms contribute to next-day learning capabilities has yet to be directly examined in empirical research.

In this pre-registered study (osf.io/78dja), we tested the hypothesis that the extent to which individuals consolidate new memories during sleep predicts their ability to encode novel information the following day, and that SWA (0.5-4 Hz) contributes to this relationship. In a within-subjects, crossover design, healthy young adults were trained on a visuospatial memory task before a night of either EEG-monitored sleep or total sleep deprivation, and were tested the following morning. Afterwards, participants were trained on a novel paired-associates task, but were not tested until 48 h later (allowing for recovery sleep in the sleep deprivation condition). Retrieval performance on the visuospatial memory and paired-associates tests thus provided independent metrics of overnight consolidation and next-day learning, respectively.

We chose these particular memory tasks because they are both reliant on hippocampus (Eichenbaum 2004; Konkel and Cohen 2009) and the Active Systems framework is primarily concerned with the overnight consolidation of hippocampus-dependent memories (Born and Wilhelm 2012; Klinzing et al. 2019; Rasch and Born 2013; Walker 2009). Moreover, previous work has consistently shown that the consolidation of both visuospatial and paired-associate memories is bolstered by overnight sleep (Ashton and Cairney 2021; Ashton et al. 2020; Cairney et al. 2018b). We reasoned that employing two conceptually different tasks was optimal as this would ensure that any potential relationship between overnight consolidation and next-day learning would not be influenced by retroactive or proactive interference.

By comparing overnight sleep and sleep deprivation, we could also investigate how protracted wakefulness affects the neural correlates of learning. Specifically, EEG recordings were acquired during paired-associates learning to test the hypothesis that sleep deprivation disrupts theta (4-8 Hz) and gamma (> 40 Hz) synchronisation, which support item binding in episodic memory (Henin et al. 2019; Köster et al. 2018; Osipova et al. 2006; Summerfield and Mangels 2005). Furthermore, in an exploratory analysis, we investigated the effect of sleep deprivation on 12-20 Hz beta desynchronization; an established marker of successful learning (Griffiths et al. 2016; Hanslmayr et al. 2014; Hanslmayr et al. 2009; Hanslmayr et al. 2012; Hanslmayr et al. 2011). Understanding how sleep disturbances impair learning and memory is increasingly important in modern society, where many people fail to regularly obtain an adequate amount of sleep (Becker et al. 2018; Bonnet and Arand 1995; Stranges et al. 2012).

## Materials and Methods

### Participants

Fifty-nine participants (32 females, mean ± SD age = 20.10 ± 1.80) were recruited on a voluntary basis and completed a preliminary session (see below). After the preliminary session, ten participants were excluded for not meeting the performance criterion and one participant was excluded for not meeting the study requirement of being a native English speaker. Among those individuals who met the performance criterion of the preliminary session, eighteen participants withdrew due to being unable to commit to the main study schedule. Our final sample size was N = 30 participants (17 females, mean ± SD age, 20.10 ± 1.65), each of whom completed both the sleep and sleep deprivation conditions (order counterbalanced, see Figure 1a). Following standard procedures in our laboratory (Ashton et al. 2019; Harrington et al. 2021a; Harrington et al. 2021b; Strachan et al. 2020), participants were asked to refrain from caffeine and alcohol for 24 h and 48 h, respectively, before each study session. Participants reported no history sleep or psychiatric disorders. Written informed consent was obtained from all participants in line with the requirements of Research Ethics Committee of the Department of Psychology at the University of York. Participants received £100 compensation upon completion of the study.

**Figure 1.**
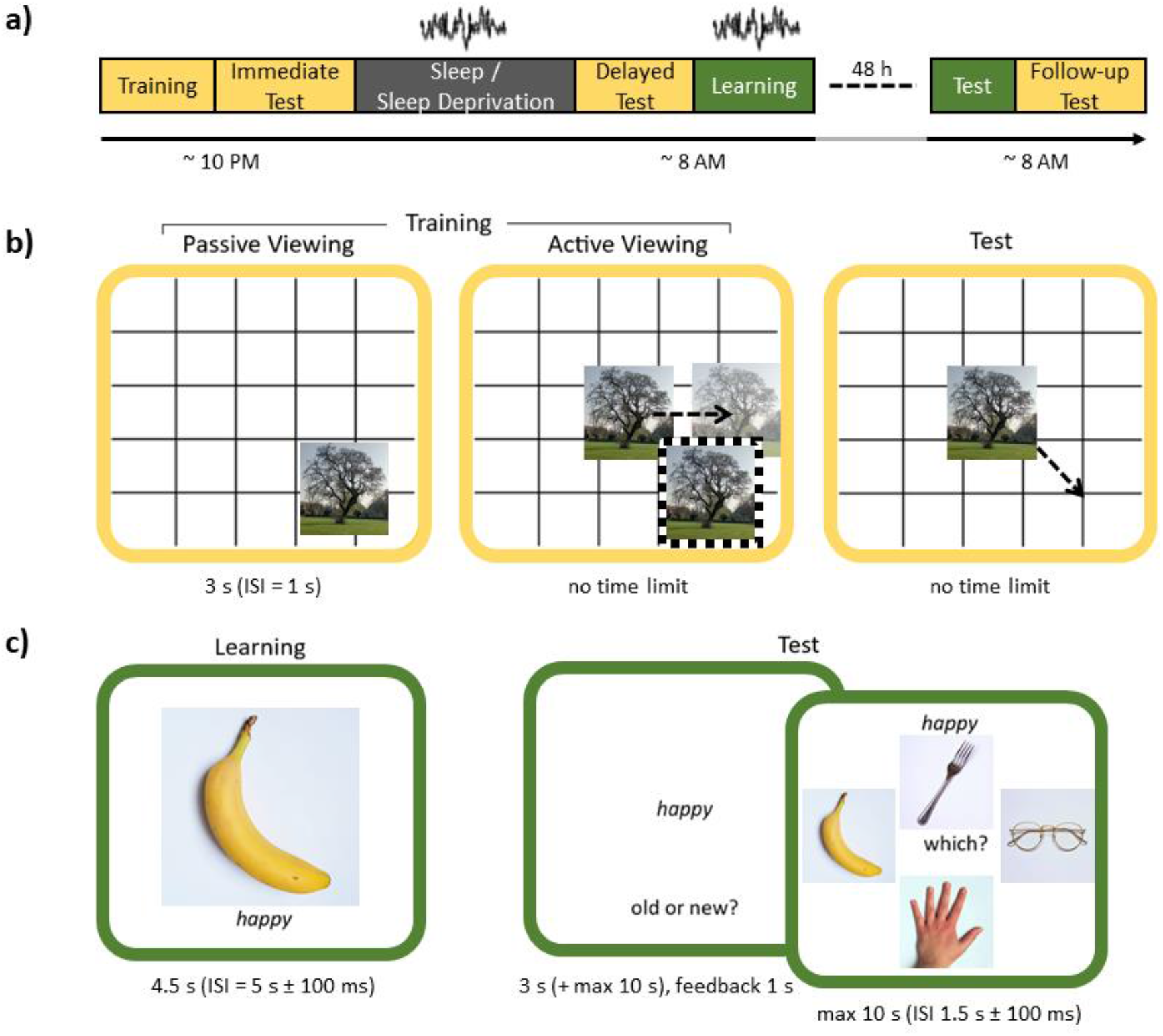
Experimental procedures and tasks. **a)** Study timeline. The colours represent the tasks: visuospatial task in yellow (see b) and paired-associates task in green (see c). Participants arrived in the evening to complete the visuospatial task (training and immediate test). After overnight sleep or sleep deprivation, participants were tested again (delayed test) and then completed the learning phase of the paired-associates task. Participants returned 48 h later (after recovery sleep) to complete the paired-associates test and a follow-up visuospatial memory test. EEG recordings were acquired during sleep and paired-associates learning. The study was a within-subjects comparison of sleep and sleep deprivation (condition order counterbalanced across participants). ISI = Interstimulus interval. **b)** Visuospatial memory task. Participants completed one round of passive viewing, during which they viewed the location of each image on a grid. Next, in the active viewing phase, participants moved each image to the location that they thought it had appeared during passive viewing and received feedback on its correct location (dashed frame). Active viewing continued until participants had met the performance criterion for all images (< 4.8cm from correct location, mean ± SD number of rounds to meet criterion: Sleep: 8.77 ± 2.39, Sleep Deprivation: 9.07 ± 2.89). The test phases followed the same procedures as one round of active viewing, but no feedback was provided. Each test trial provided an accuracy score in cm, which described how far the image was placed from its correct location. **c)** Paired-associates task. Participants completed one round of learning, during which they encoded adjective-image pairings. At test, a word was presented in isolation and participants first indicated whether it was ‘old’ (i.e. they recognised the word from learning) or ‘new’ (i.e. they did not recognise the word from learning). For correctly recognised (old) words, participants then indicated which of the four presented images was associated with that word at learning. For words identified as new, or for previously unseen words that were incorrectly identified as old, participants moved immediately onto the next trial.

Statistical power was calculated prior to data collection using an effect size of d = 0.56 from Ashton et al. (2020). This effect size was derived from a paired-samples t-test comparing forgetting after a night of sleep or total sleep deprivation. Based on this effect size, our pre-registered sample of N = 30 participants provided 83.7% power (alpha = .05, two-tailed).

### Tasks and Stimuli

#### Visuospatial task (see Figure 1b)

One-hundred images of neutral scenes were taken from the International Affective Picture System (Lang et al. 1997) and the Nencki Affective Picture System (Marchewka et al. 2014). These were divided into two sets of 50 images for use in the sleep and sleep deprivation conditions (assignment of image set to condition was counterbalanced). The visuospatial task was divided into three phases:

##### 1. Training I: Passive viewing

Each of the 50 images was presented in a randomly selected location on a grid background (exposure time = 3 s, interstimulus interval [ISI] = 1 s). Participants were instructed to try and memorise the image locations for a later test. Image presentation order was randomised.

##### 2. Training II: Active viewing

Each image appeared in the centre of the grid and participants moved it to the location that they believed it had appeared at passive viewing. The image then reappeared in its correct location to serve as feedback. This continued until all images had been placed < 4.8 cm (< 150 pixels) from their correct location on two consecutive rounds of active viewing (images for which this criterion was met were dropped from subsequent active viewing rounds). Image presentation order was randomised.

##### 3. Test

The test phase followed the same procedures as one round of active viewing with the exception that no feedback was provided. Three tests were completed (immediate, delayed and follow-up).

#### Paired-associates task (see Figure 1c)

Two hundred images of natural and manmade objects on a white background were taken from Konkle et al. (2010) and online resources. These were divided into two sets of 100 objects (50 natural and 50 manmade) for use in the sleep and sleep deprivation conditions (assignment of object set to condition was counterbalanced). Three hundred adjectives (150 adjectives per condition, assignment counterbalanced) were taken from Cairney et al. (2018a). Within each condition, 100 adjectives were randomly selected as targets and the remaining 50 as foils.

##### 1. Adjective familiarisation

Each of the 100 target adjectives was presented for 3 s. Participants were instructed to rate how often they would use each adjective in conversation on a scale of 1 to 9 (1 = never, 5 = sometimes and 9 = often) within an additional 4 s (ISI with fixation crosshair = 1.5 s ± 100ms). Adjective presentation order was randomised.

##### 2. Image familiarisation

Each of the 100 images (50 natural and 50 manmade objects) was presented for 3 s. Participants were instructed to imagine themselves interacting with each object and then categorise it as being natural or manmade within an additional 4 s (ISI with fixation crosshair = 1.5 s ± 100ms). Image presentation order was randomised.

##### 3. Learning

On each trial, participants were presented with an adjective and image from each of the prior familiarisation phases for 4.5 s and instructed to memorise the adjective-image pairing for a future test. To facilitate learning, participants were asked to create a story or mental image in their mind that involved the adjective and image interacting for the full duration of the trial, and then to rate this association as realistic or bizarre within an additional 4 s. A longer ISI of 5 s (± 100ms) was used to facilitate the analysis of EEG data acquired during adjective-image learning (this comprised a 2 s progression bar followed by 3 s of fixation). Adjective-image pairing order was randomised.

##### 4. Test

Each of 150 adjectives (100 from learning and 50 unseen foils) was presented for 3 s. Participants were first instructed to indicate whether the adjective was old or new within an additional 10 s. Feedback on accuracy (correct/incorrect) was then provided for 1 s. For correct old responses, participants were presented with four images (all of which had been seen at learning) and asked to indicate which image was paired with the adjective within 10 s. Participants then rated how confident they were in their response on a scale of 1 (not confident) to 4 (very confident) within 10 s. For incorrect old responses or new responses, participants moved immediately onto the next trial (ISI with fixation crosshair = 1.5 s ± 100 ms). Adjective presentation order was randomised.

#### Psychomotor vigilance task (PVT)

The PVT was obtained from Khitrov et al. (2014, bhsai.org/downloads/pc-pvt). Participants were instructed to respond when a digital counter appeared on the screen (ISI = 2-10 s). Participants received feedback on their response times and the task lasted for 3 min.

### Procedure

#### Preliminary session

Participants completed a preliminary memory assessment before entering the main study. They learned 180 semantically related word pairs (e.g. *Horizon* – *Sun*) and were immediately tested with a cued recall procedure. Participants scoring between 50% and 95% were invited back for the main experiment. This ensured that participants were unlikely to perform at floor or ceiling in the visuospatial and paired-associates tests of the main study.

#### Session one: evening

Participants arrived between 8:30 PM and 10 PM. In the sleep condition (earlier arrivals), participants were immediately wired-up for overnight EEG monitoring. Participants began the study by completing the Stanford Sleepiness Scale (Hoddes et al. 1973), PVT and then the training and immediate test phases of the visuospatial task.

#### Overnight interval

In the sleep condition, participants went to bed at ∼11 PM and were woken up in the morning at ∼7 AM (thus achieving ∼8 h of EEG-monitored sleep). In the sleep deprivation condition, participants remained awake across the entire night under the supervision of a researcher. During the sleep deprivation period, participants were provided with refreshments and were permitted to play games, watch movies or complete coursework. Because our sample was mostly made up of university students and a significant number of daytime study hours would be lost as a result of overnight sleep deprivation, we chose to allow participants to complete coursework in order to facilitate recruitment. Importantly, all of the permitted activities were deemed suitable because they were not conceptually linked to the materials that participants had learned the previous evening (i.e. object-location associations) or would learn the following morning (i.e. adjective-image pairings).

#### Session two: morning

Participants in the sleep deprivation condition were wired-up for EEG monitoring (this was not necessary in the sleep condition as electrodes had already been attached the previous night). Participants then completed another round of the Stanford Sleepiness Scale and PVT, and another (delayed) visuospatial test. Afterwards, participants carried out the familiarisation phases of the paired-associates task, before completing the paired-associates learning phase with EEG monitoring. Participants were not given any explicit instruction on what to do (e.g. when to go to sleep) during the 48-h interval that preceded session three.

#### Session three: follow-up

Participants returned 48 h after session two (thereby allowing for recovery sleep in the sleep deprivation condition) and completed a final round of the Stanford Sleepiness Scale and the PVT. They then carried out the paired-associates test and a final (follow-up) visuospatial test.

### Equipment

#### Experimental tasks

All tasks were executed on a Windows PC and participant responses were recorded with a keyboard or mouse. The visuospatial task was implemented in Presentation version 14.1 (Neurobehavioural Systems, Inc.) and the paired-associates task was implemented in Psychtoolbox 3.0.13 (Brainard 1997; Kleiner et al. 2007; Pelli 1997) and MATLAB 2019a (The MathWorks, Inc.).

#### EEG

EEG recordings were administered with two Embla N7000 systems and one Embla NDx system with REMLogic 3.4 software. The Embla NDx was acquired when upgrading our sleep laboratory from a two- to three-bedroom facility (the N7000 was no longer available for purchase). For all but three participants, the same EEG system was used in the sleep and sleep deprivation conditions. Gold-plated electrodes were attached to the scalp according to the international 10-20 system at frontal (F3 and F4), central (C3 and C4), parietal (P3 and P4) and occipital (O1 and O2) locations, and referenced to the linked mastoids. Left and right electrooculography electrodes were attached, as were electromyography electrodes at the mentalis and submentalis bilaterally, and a ground electrode was attached to the forehead. An additional reference electrode was placed at Cz for the NDx system. We ensured that all electrodes had a connection impedance of < 5 kΩ immediately before any EEG data was collected (i.e. for participants in the sleep condition, impedances were checked before sleep and again in the morning before the learning task). Any electrodes that fell above this threshold were replaced and re-checked. All online signals were digitally sampled at 200 Hz (N7000) or 256 Hz (NDx, down-sampled to 200 Hz during preprocessing).

#### Actigraphy

Participants wore wristwatch actigraphy devices (Actiwatch 2, Philips Respironics, USA). throughout the study so that we could monitor their sleep when they were outside of the laboratory.

### Data analyses

#### Behaviour

Behavioural data were analysed using R Studio (v.1.4.1717, RStudio Team 2021). Memory consolidation was indexed by the change in visuospatial memory accuracy between the immediate and delayed test. For each participant and test, we computed an error score for each image by calculating the distance (cm) between the recalled location (image centre) and the location that the image had appeared at passive viewing. We derived a retention index (RI) by subtracting the error score at the delayed test from the error score at the immediate test for each image, and then averaging across images. A follow-up RI was calculated between the immediate and follow-up tests using the same method. To ease understanding (e.g. higher RI = better retention), we swapped the order of the RI subtraction to that which was pre-registered. This change yields statistically identical results aside from the sign change. As pre-registered, one participant was removed from analyses that included RI^SleepBenefit^ scores (see below) because their RI at the delayed test in the sleep deprivation condition was > 3 SD from the mean.

Next-day learning was assessed by the learning index (LI), which equated to the percentage of correctly recognised images on the paired-associates test. Between-condition differences in RI and LI were analysed using paired-samples t-tests with a significance threshold of p < .05. We report the “classical” Cohen’s d as our effect size estimate because it is unaffected by experimental design and thus facilitates comparisons across different studies (R function: cohensD, R package: lsr, Navarro 2015).

One of our primary aims was to investigate the relationship between sleep-associated consolidation and next-day learning, and how SWA contributes to this relationship. To do this, we first quantified the benefit of sleep (vs sleep deprivation) on the RI and LI. We subtracted (for each participant) the RI in the sleep deprivation condition from the RI in the sleep condition to derive a RI^SleepBenefit^. Similarly, we subtracted (for each participant) the LI in the sleep deprivation condition from the LI in the sleep condition to obtain a LI^SleepBenefit^. Positive scores on the RI^SleepBenefit^ and LI^SleepBenefit^ therefore indicate a sleep-associated improvement in performance. RI^SleepBenefit^ and SWA (see below) were entered as predictors of LI^SleepBenefit^ in a forced-entry multiple regression analysis. A Bayesian multiple regression analysis (R package: BayesFactor, Morey and Rouder 2018) was used to test for evidence of the null (i.e. no relationship between sleep-associated consolidation [RI^SleepBenefit^], SWA and next-day learning [LI^SleepBenefit^]). Exploratory correlations were computed using Pearson’s R.

#### EEG (Sleep)

##### Preprocessing

Sleep EEG data were partitioned into 30 s epochs and scored in RemLogic 3.4 according to standardised criteria (Iber 2007). Epochs scored as sleep stage N2 or slow-wave sleep (SWS) were exported to MATLAB 2019a using the FieldTrip toolbox (Oostenveld et al. 2011, v.10/04/18) for further analysis. Artifacts were identified and removed using FieldTrip’s Databrowser (mean ± SD artifacts rejected, 3.5 ± 2.85), noisy channels were removed (four channels across four participants) and two entire datasets were excluded due to excessive noise. The remaining data were band-pass filtered between 0.3 and 30 Hz using Butterworth low-pass and high-pass filters.

##### Power spectral analysis

Due to significant noise in the occipital channels (as a result of electrodes detaching during the night in several participants), we only included frontal (F3 and F4), central (C3 and C4) and parietal (P3 and P4) channels in our spectral analysis of the sleep EEG data. Using functions from the FieldTrip toolbox, artifact-free N2 and SWS epochs were applied to a Fast Fourier Transformation with a 10.24 s Hanning window and 50% overlap. EEG Power in the SWA (0.5-4 Hz) and fast spindle (12.1-16 Hz) bands was determined by averaging across the corresponding frequency bins and across channels.

#### EEG (Learning)

##### Preprocessing

All eight EEG channels (F3, F4, C3, C4, P3, P4, O1 and O2) were included in our analysis of learning. Data were high-pass filtered (0.5 Hz), notch filtered (49-51 Hz), and segmented into trials (−3 s to 4.5 s around stimulus onset). Trials for which participants did not provide a response were removed from the analysis (mean ± SD excluded trials, sleep: 0.1 ± 0.45, sleep deprivation: 5.11 ± 7.93, Priest et al. 2001). From scalp electrodes, eye-blinks and cardiac components were identified and removed using an independent components analysis, and noisy channels were interpolated via a weighted-average of their nearest neighbours (fourteen channels across six participants and two conditions). Trials were visually inspected and data from two participants were removed due to excessive noise in multiple channels.

##### Time-frequency analyses

Time-frequency representations (TFRs) were calculated separately for lower (4-30 Hz) and higher frequencies (30-60 Hz). Our pre-registered upper bound was 120 Hz, but because our sampling rate was 200 Hz the upper bound was above the Nyquist frequency and had to be lowered. For lower frequencies, data were convolved with a 5-cycle Hanning taper in 0.5 Hz frequency steps and 5 ms time steps using an adaptive window-length (i.e. where window length decreases with increasing frequency, e.g. 1.25 s at 4 Hz and 1 s at 5 Hz, to retain 5 cycles). For higher frequencies, data were convolved with tapers of Slepian sequence (3 tapers), also in steps of 0.5 Hz and 5 ms with an adaptive window-length. For this latter analysis, frequency smoothing was set to 0.4 of the frequency of interest (e.g. 20 Hz smoothing at 50 Hz). Artifact rejection was achieved via a data-driven approach, applied separately to the analyses of lower and higher frequencies: power values that exceeded the 85^th^ percentile across all time/frequency points and trials were removed from each participant’s dataset. TFRs were converted into percent power change relative to a -400 to -200 ms pre-stimulus baseline window. This window was chosen to mitigate baseline contamination by post-stimulus activity while preserving proximity to stimulus onset (note that our post-stimulus time-window of interest started at 0.3 s, see below). Trials were divided into subsequently remembered and forgotten adjective-image pairings (based on the test phase 48 h later). Because our 49-51 Hz notch filter overlapped with our gamma frequency range, we re-ran our higher frequency analysis (30-60 Hz) without a notch filter and the results in the gamma frequency range (40-60 Hz) were unchanged.

##### Statistics

TFR analyses were performed as dependent samples analyses and corrected for multiple comparisons using FieldTrip’s nonparametric cluster-based permutation method (1000 randomisations). Clusters were defined by channel * time whilst averaging across the frequency bands of interest (theta [4-8 Hz], alpha [8-12 Hz], beta [12-20 Hz] and gamma [40-60 Hz], cluster threshold p < .05). Pre-registered analyses in theta and gamma bands were one-tailed, whereas exploratory analyses were two-tailed. To reduce interference from early visual evoked responses, the time window of interest was set from 0.3-2 s (Friedman and Johnson 2000; Osipova et al. 2006). A factorial approach was used to assess the impacts of sleep deprivation (vs sleep) on the neural correlates of encoding: we calculated the grand average TFR difference for subsequently remembered > forgotten adjective-image pairings within each condition (sleep and sleep deprivation), and then entered these contrasts into the cluster-based permutation analysis (Sleep^remembered>forgotten^ > Sleep Deprivation^remembered>forgotten^). To reflect the rationale of the cluster-based permutation test, we report effect sizes as Cohen’s d_z_ based on the average of the largest cluster (i.e. averaging across all channels and time points that contributed at any point to the largest cluster, Meyer et al. 2021).

## Results

### Sleep benefits memory consolidation

To assess overnight consolidation, we computed a retention index (RI) from the immediate and delayed visuospatial memory tests (higher RI = better overnight retention, see Materials and Methods). As expected, the RI was significantly higher after sleep than sleep deprivation (t(28) = 3.78, p < .001, d = 0.71, see Figure 2a). To ensure that our findings were not driven by between-condition differences in fatigue at the delayed test, we also assessed memory retention between the immediate and follow-up test (which took place 48 h after the delayed test, thereby allowing for recovery sleep). As expected, the RI was still significantly higher in the sleep (vs sleep deprivation) condition (t(28) = 2.18, p = .038, d = 0.44, see Figure 2b), suggesting that sleep had facilitated overnight consolidation. There was no significant between-condition difference in visuospatial accuracy at the immediate test (mean ± SEM, sleep: 2.44 ± 0.10, sleep deprivation: 2.57 ± 0.10, t(28) = -0.98, p = .337, d = 0.19, BF_01_ = 3.28), and no difference in the benefit of sleep on retention (RI^SleepBenefit^ [i.e. sleep condition RI – sleep deprivation condition RI], see below) between participants who completed the sleep condition before or after the sleep deprivation condition (t(27) = 0.22, p = .828, d = 0.08, BF_01_ = 2.81).

**Figure 2.**
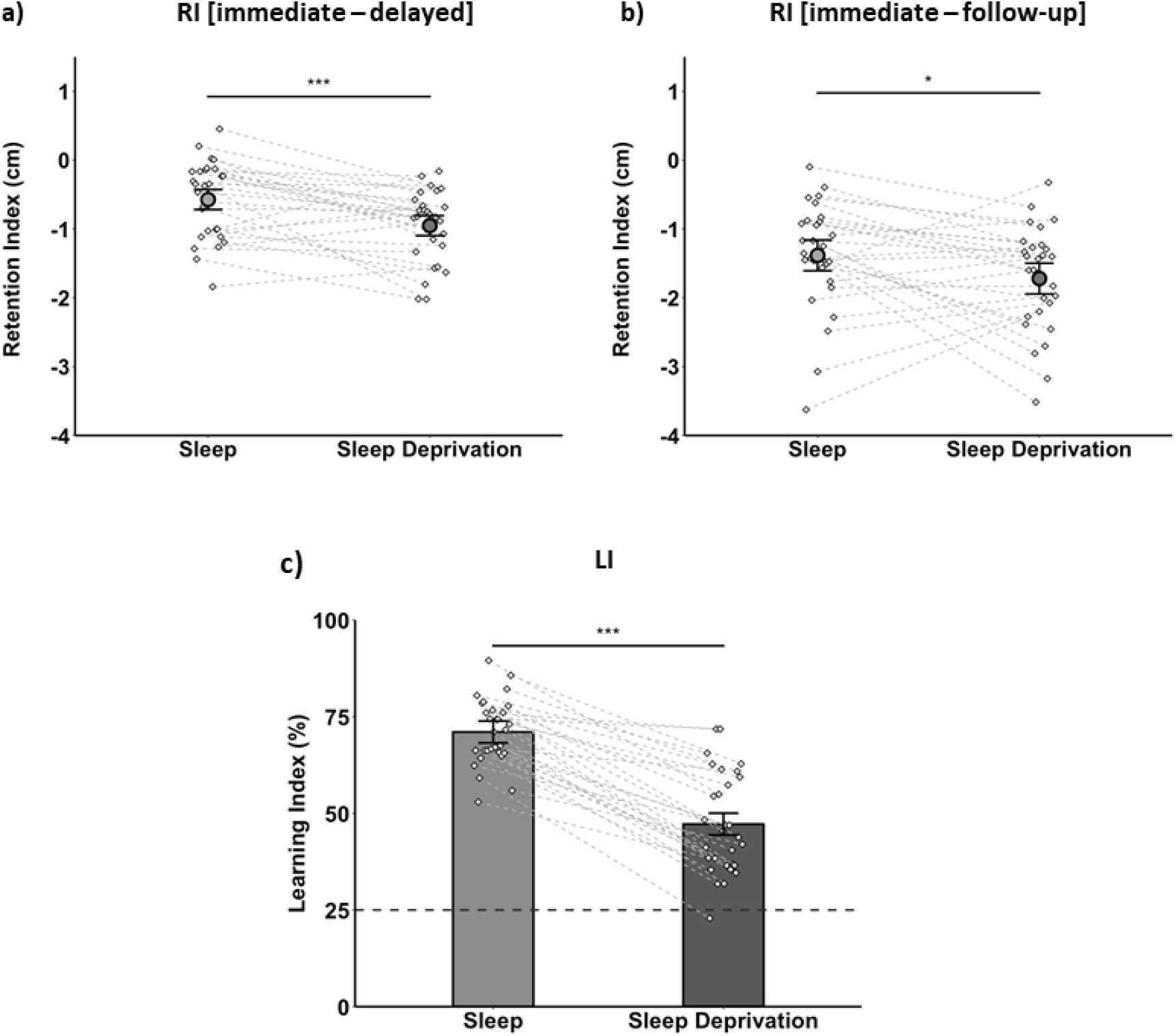
Behaviour. **a)** Retention index (RI) between the immediate and delayed test, and **b)** between the immediate and follow-up test (i.e. 48 h after the delayed test, following recovery sleep). A higher RI indicates better retention. **c)** Learning index (LI, tested 48 h after sleep or sleep deprivation). Higher scores indicate better learning and the dashed line represents chance performance (25%). All figures show condition means (± SEM). Diamonds and connecting lines represent individual participants. *** p < .001, * p < .05.

Although response times on the PVT were slower in the morning after sleep deprivation (mean ± SEM, 399.00ms ± 17.63) than sleep (289.15 ± 4.34, p < .001), there was no significant relationship between RI^SleepBenefit^ and PVT^SleepBenefit^ (i.e. [mean RT after sleep – mean RT after sleep deprivation], R^2^ = -.15, p = .440, BF_01_ = 1.92). Similarly, as indicated by the Stanford Sleepiness Scale (SSS), participants felt less alert after sleep deprivation (mean ± SEM, 5.37 ± 0.15) than sleep (2.27 ± 0.16). However, there was no significant correlation between RI^SleepBenefit^ and SSS^SleepBenefit^ (i.e. [mean rating after sleep – mean rating after sleep deprivation], R^2^ < -.01, p = .991, BF_01_ = 2.46). Extended analysis of the PVT and Stanford Sleepiness Scale data is available in the Supplementary Material.

### Sleep improves next-day learning

To assess encoding performance after sleep or sleep deprivation, we calculated a learning index (LI), which equated to the percentage of correctly recognised images on the paired-associates test (this took place 48 h after encoding, following recovery sleep). As expected, encoding performance was significantly higher after sleep than sleep deprivation (t(29) = 12.19, p < .001, d = 2.17, see Figure 2c), suggesting that sleep had benefited next-day learning. There was no significant difference in the benefit of sleep on new learning (LI^SleepBenefit^ [i.e. sleep condition LI – sleep deprivation condition LI], see below) between participants who completed the sleep condition before or after the sleep deprivation condition (t(28) = 0.37, p = .712, d = 0.14, BF_01_ = 2.75).

There was no significant relationship between LI^SleepBenefit^ and PVT^SleepBenefit^ (R^2^ = -.30, p = .113), although the evidence for the null remained inconclusive (BF_01_ = 0.86). Similarly, there was no significant correlation between LI^SleepBenefit^ and SSS^SleepBenefit^ (R^2^ = -.35, p = .056) with inconclusive evidence for the null (BF_01_ = 0.53).

### No relationship between sleep-associated consolidation, slow wave activity and next-day learning

Next, we tested the hypothesis that overnight consolidation predicts next-day learning, and that slow-wave activity (SWA) contributes to this relationship. Because our aim was to target the relationship between sleep-associated memory processing and next-day learning, it was necessary to first quantify the positive impact of sleep (vs sleep deprivation) on the RI and LI. We therefore subtracted both the RI and LI between the sleep and sleep deprivation conditions (on a participant-by-participant basis), such that positive scores on the resultant RI^SleepBenefit^ and LI^SleepBenefit^ metrics indicated a sleep-associated improvement in performance. SWA was defined as EEG power within the 0.5-4 Hz frequency band during sleep stages N2 and slow-wave sleep (SWS, collapsed across all EEG channels). In a multiple regression model, we entered RI^SleepBenefit^ and SWA as predictors and LI^SleepBenefit^ as the outcome variable. Contrary to expectations, sleep-associated consolidation (RI^SleepBenefit^) and SWA did not significantly account for next-day learning (LI^SleepBenefit^, F(2,24) = 1.51, R^2^ = 0.11, p = .242, see Figure 3). No significant relationship was observed between RI^SleepBenefit^ and LI^SleepBenefit^ independently of SWA (B = 3.30, t(24) = 0.86, p = .399), nor between SWA and LI^SleepBenefit^ independently of RI^SleepBenefit^ (B = -0.51, t(24) = -1.65, p = .111). RI^SleepBenefit^ did not significantly correlate with SWA (R^2^ = 0.21, p = .298). A follow-up Bayesian analysis revealed anecdotal evidence in support of the null (i.e. that sleep-associated consolidation and SWA did not account for next-day learning, BF_01_ = 2.04).

**Figure 3.**
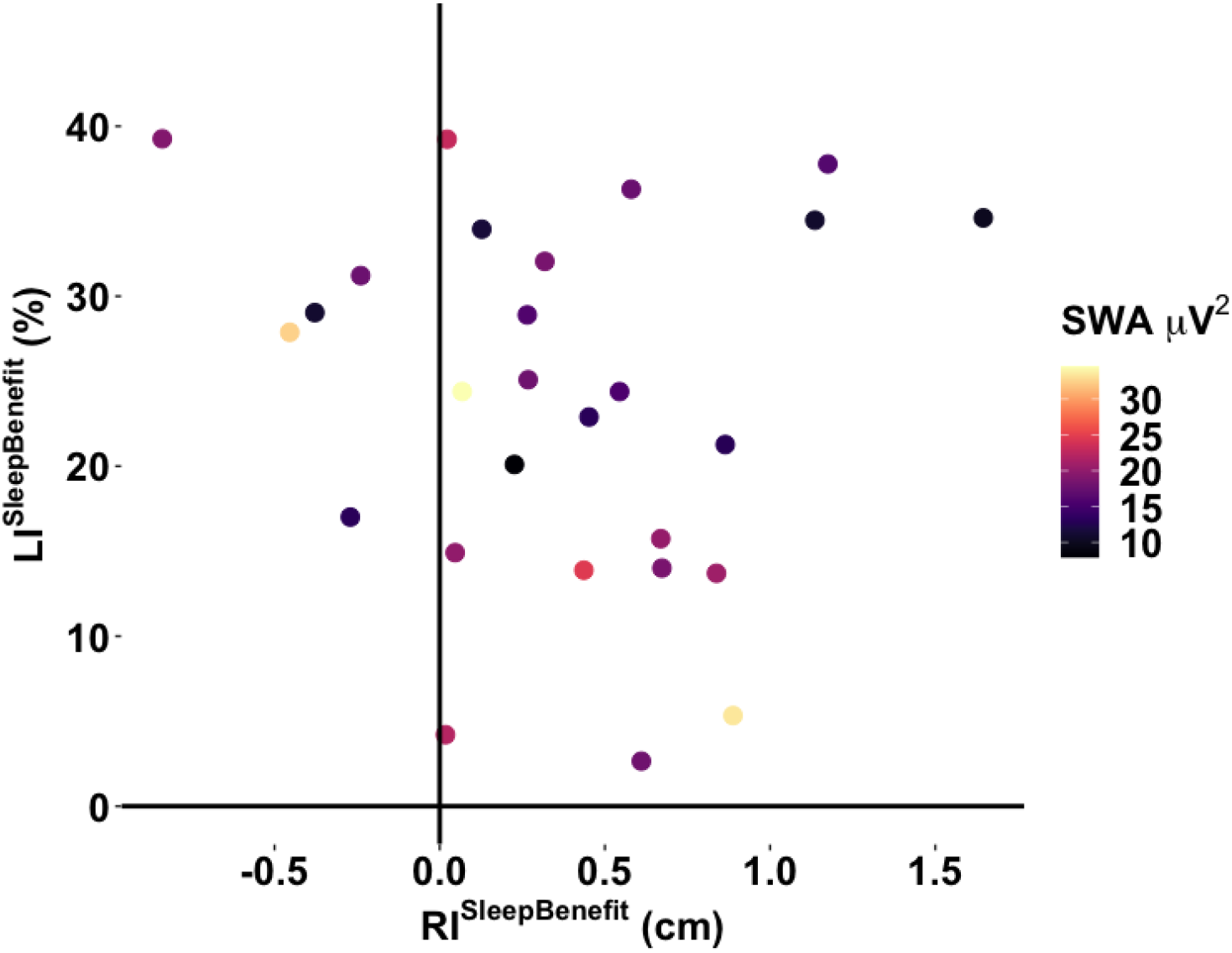
Relationship between sleep-associated consolidation, SWA and next-day learning. Sleep-associated consolidation (RI^SleepBenefit^) and SWA had no significant impact on next-day learning (LI^SleepBenefit^).

In a subsidiary analysis, we repeated this multiple regression but only entered data from the sleep condition into our model (i.e. the RI^SleepBenefit^ and LI^SleepBenefit^ were replaced with the RI and LI from the sleep condition alone). Our findings mirrored those of the foregoing analysis: sleep-associated consolidation (RI) and SWA did not significantly account for next-day learning (LI, F(2,25) = 1.83, p = .181, R^2^ = 0.13, BF_01_ = 1.68). There was also no significant relationship between RI and LI independently of SWA (B = 4.46, t(25) = 1.67, p = .107), nor between SWA and LI independently of RI (B = -0.25, t(25) = -1.16, p = .256), and no significant correlation was observed between RI and SWA (R^2^ = 0.28, p = .143).

We also explored whether RI in the sleep condition was correlated with other sleep parameters previously implicated in declarative memory consolidation: time (min) in SWS (Backhaus et al. 2006; Scullin 2013) and fast spindle power (12.1-16 Hz, Cox et al. 2012; Tamminen et al. 2010). However, no significant relationships emerged (all p > .368). Sleep data is available in Table 1.

**Table 1:**
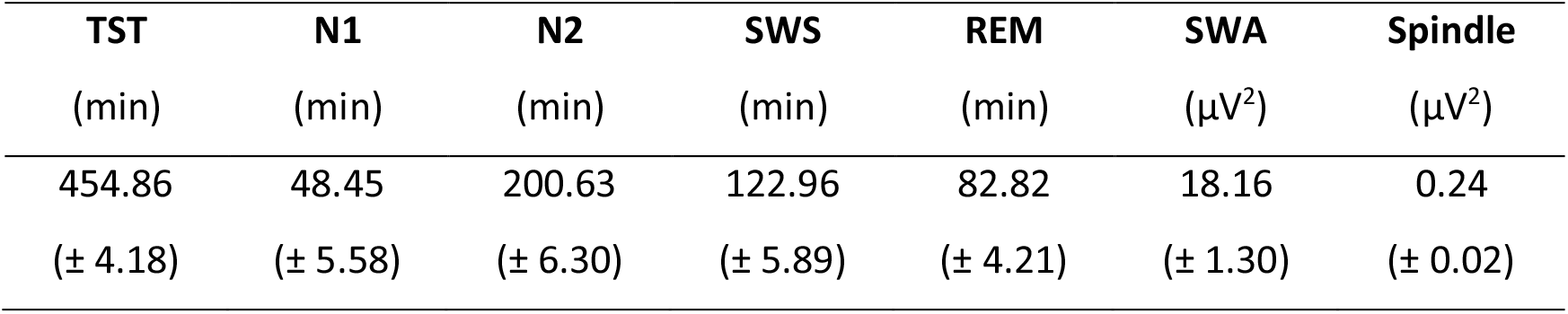
Sleep data. Parameters include total sleep time (TST), time spent in each stage of sleep (N1, N2, slow-wave sleep [SWS] and rapid eye movement sleep [REM]) and power in the slow-wave activity (SWA, 0.5-4 Hz) and fast spindle (12.1-16 Hz) bands within N2 and SWS. Data are shown as mean (± SEM).

### Sleep deprivation disrupts beta desynchronization during successful learning

Finally, we tested the hypothesis that sleep deprivation disrupts theta and gamma synchronisation at learning. However, no significant differences were observed in the theta (4-8 Hz) or gamma (40-60 Hz) bands when comparing time-frequency representations between subsequently remembered and forgotten adjective-image pairings or between the sleep and sleep deprivation conditions, and there was no significant interaction between these contrasts (all p > .05).

In an exploratory analysis, we investigated the effect of sleep deprivation on beta desynchronization; an established marker of successful learning (Griffiths et al. 2016; Hanslmayr et al. 2014; Hanslmayr et al. 2009; Hanslmayr et al. 2012; Hanslmayr et al. 2011). Consistent with these previous findings, an overall reduction in beta power was observed during encoding of subsequently remembered (vs forgotten) adjective-image pairings when combining the sleep and sleep deprivation conditions (corresponding to two clusters in the left hemisphere beginning at ∼1.5-1.7 s (p = .044, d = -.66) and ∼1.75-1.9 s (p = .038, d = -.49) after stimulus onset (see Figure 4a).

**Figure 4.**
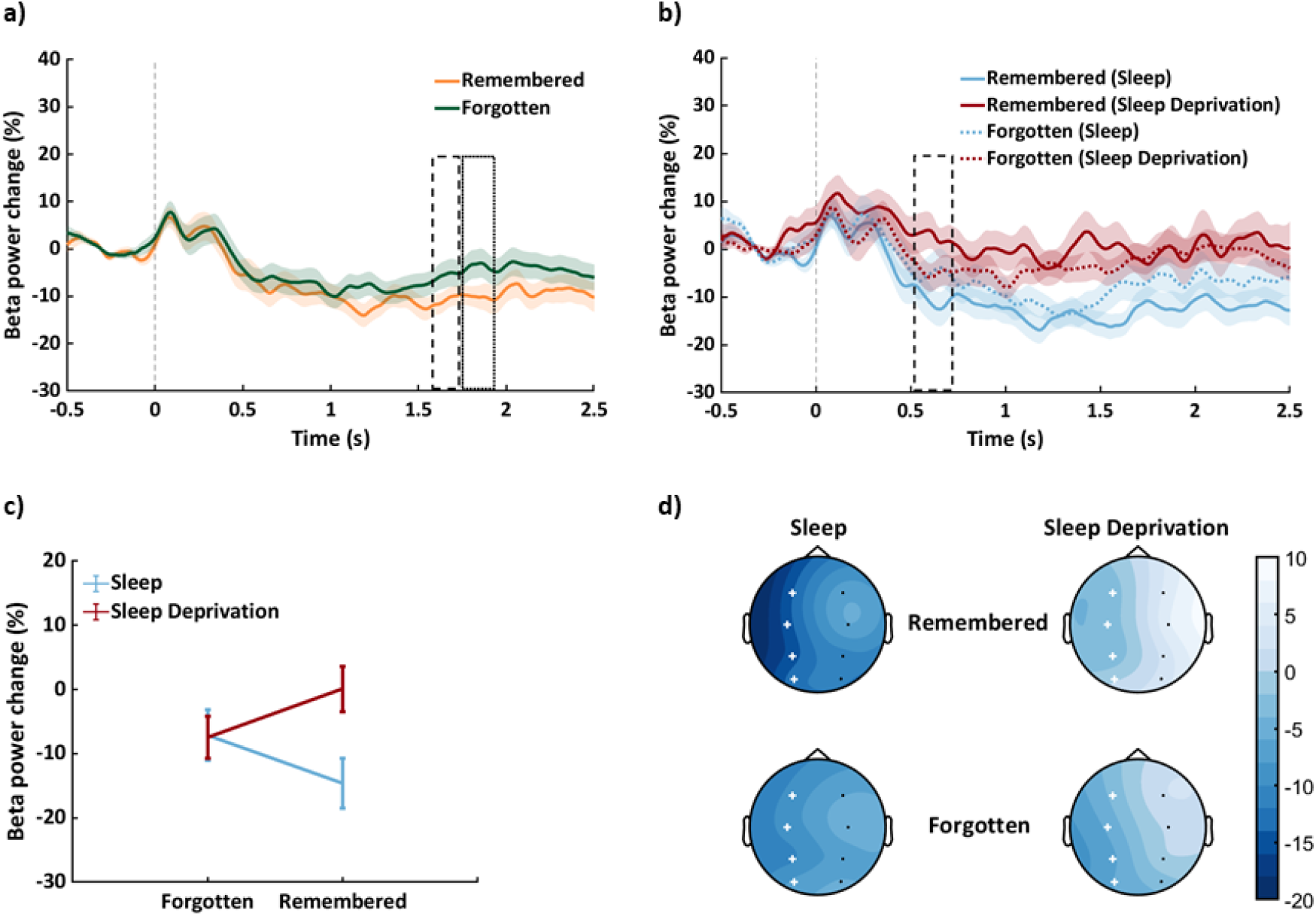
Changes in beta power during successful learning after sleep and sleep deprivation. **a)** Grand average change in beta power from baseline (12-20 Hz, all channels) for subsequently remembered and forgotten adjective-image pairings. Dashed and dotted boxes approximate the time windows contributing to the significant clusters and shaded areas represent SEM. **b)** The same contrast as in panel a, but presented separately for the sleep and sleep deprivation conditions. **c)** Corresponds to the significant cluster (interaction) in panel b, averaged across time (error bars represent SEM). Whereas encoding of subsequently remembered (vs forgotten) adjective-image pairings was associated with reduced beta power after sleep, no significant difference in beta power emerged from the same contrast after sleep deprivation. **d)** Topographical representations of the change in beta power for subsequently remembered and forgotten adjective-image pairings in the sleep and sleep deprivation conditions, averaged across the time-window of interest (0.3-2 s). Crosses represent the channels of the cluster that corresponds to the significant interaction. The vertical bar represents the change in beta power from baseline (%).

Interestingly, changes in beta power accompanying successful learning were significantly different in the sleep and sleep deprivation conditions (interaction, corresponding to a cluster in the left hemisphere at ∼0.5-0.7 s, p = .014, d = -.33, see Figures 4b and 4d). Whereas encoding of subsequently remembered (vs forgotten) adjective-image pairings was associated with a downregulation of beta power after sleep (p = .005), an apparent upregulation of beta power emerged from the same contrast after sleep deprivation (p = .019, see Figure 4c, although this latter post-hoc test did not survive Bonferroni correction, alpha = .0125). Moreover, beta power was significantly reduced during encoding of subsequently remembered pairings in the sleep (vs sleep deprivation) condition (p = .001), but no such difference was observed during encoding of subsequently forgotten pairings (p = .928).

To explore whether this significant interaction was driven by increased fatigue in the sleep deprivation (vs sleep) condition, we correlated (for each participant) average beta power for the contrast Sleep^remembered>forgotten^ > Sleep Deprivation^remembered>forgotten^ (within the significant group-level cluster) with SSS^SleepBenefit^ and PVT^SleepBenefit^. No significant relationships were observed (SSS: R^2^ = -0.20, p = .311, BF_01_ = 1.58, PVT: R^2^ = .18, p = .371, BF_01_ = 1.73), suggesting that the foregoing findings did not arise from between-condition differences in fatigue.

An overall reduction in beta power was also observed for the sleep (vs sleep deprivation) condition when combining subsequently remembered and forgotten adjective-image pairings, corresponding to two clusters in the left (∼1-1.5 s, p = .014, d = -0.63) and right hemisphere (∼1.3-1.7 s, p = .038, d = -0.47).

Given the previously reported links between alpha (8-12 Hz) desynchronization and successful learning (Griffiths et al. 2016; Weisz et al. 2020), we also explored activity in this frequency band (same contrasts as above), but no significant effects were observed (all p > .05).

### Actigraphy

Hours slept during the 48-h interval between the delayed and follow-up tests (as estimated via wristwatch actigraphy) were applied to a 2 (Condition: Sleep/Sleep Deprivation) * 2 (Night: One/Two) repeated measures ANOVA (R function: anova_test, R package: rstatix). Note that two participants were not included in this analysis due to technical problems with the actigraphy device. There was a main effect of Night (F(1,27) = 62.47, p < .001, η_p_^2^ = .70), indicating that all participants slept for longer on night one than night two. A significant Condition * Night interaction (F(1,27) = 14.21, p < .001, η_p_^2^ = .35) also emerged, with Bonferroni-corrected post-hoc tests indicating that sleep duration was longer in the sleep deprivation (vs sleep) condition on night one (mean ± SEM hours sleep, sleep deprivation: 9.03 ± 0.45, sleep: 7.52 ± 0.24, p = .006) but shorter on night two (sleep deprivation: 5.33 ± 0.23, sleep: 6.00 ± 0.25, p = .036). There was no main effect of Condition (F(1,27) = 3.07, p = 091, η_p_^2^ = .10).

It is possible that the longer duration of sleep on the first night after learning in the sleep deprivation (vs sleep) condition augmented the consolidation of newly learned adjective-image pairings, potentially mitigating the initial impact of sleep loss on encoding. To test this possibility, we correlated the between-condition difference in sleep duration on the first night after learning (sleep deprivation condition – sleep condition) with the LI^SleepBenefit^. However, no significant relationship emerged (R^2^ = -0.06, p = .756, BF_01_ = 2.33).

## Discussion

Sleep provides a benefit over wake for retaining memories and also for learning new ones (Alberca-Reina et al. 2014; Ashton and Cairney 2021; Ashton et al. 2020; Cairney et al. 2018b; Cousins et al. 2018; Durrant et al. 2016; Gais et al. 2006; Gaskell et al. 2018; Kaida et al. 2015; Payne et al. 2012; Talamini et al. 2008; Tempesta et al. 2016; Yoo et al. 2007). Some suggest that these benefits can be explained by an active role of slow-wave sleep (SWS) and associated neural oscillations in shifting the memory retrieval network from hippocampus to neocortex, and thus restoring hippocampal capacity for new learning (Born and Wilhelm 2012; Klinzing et al. 2019; Rasch and Born 2013; Walker 2009). In the current study, we tested the hypothesis that the extent to which individuals consolidate new memories during sleep predicts their ability to encode new information the following day, and that slow-wave activity (SWA) contributes to this relationship. Although we observed a benefit of sleep (relative to sleep deprivation) on our measures of overnight consolidation and next-day learning, we found no evidence of a relationship between the two measures, nor with SWA.

Given the importance of sleep for new learning, we further sought to understand how sleep deprivation affects the neural correlates of successful encoding. Interestingly, whereas learning of subsequently remembered (vs forgotten) associations was associated with a downregulation of 12-20 Hz beta power after sleep (as reported in previous work, Griffiths et al. 2019; Hanslmayr et al. 2014; Hanslmayr et al. 2009; Hanslmayr et al. 2012; Hanslmayr et al. 2011), no significant difference in beta power emerged after sleep deprivation. These findings suggest that an absence of sleep disrupts the neural operations underpinning memory encoding, leading to suboptimal performance.

### Sleep benefits overnight consolidation and next-day learning

Previous work has shown that sleep supports memory consolidation (Ashton and Cairney 2021; Ashton et al. 2020; Cairney et al. 2018b; Durrant et al. 2016; Gais et al. 2006; Gaskell et al. 2018; Payne et al. 2012; Talamini et al. 2008) and subsequent learning (Cousins et al. 2018; Kaida et al. 2015; McDermott et al. 2003; Tempesta et al. 2016; Yoo et al. 2007). In keeping with these studies, we found that memory retention and next-day learning benefited from overnight sleep relative to sleep deprivation.

Although this was a pre-registered investigation of sleep’s role in learning and memory, and was motivated by prior work on the same topic (Gais et al. 2006; Yoo et al. 2007), it is important to consider the extent to which our findings can disentangle the memory effects of sleep from the disruptive influences of sleep deprivation. Extended periods of wakefulness give rise to various cognitive impairments (Krause et al. 2017), meaning that poorer performance in the sleep deprivation (vs sleep) condition could reflect the indirect consequences of sleep loss, rather than a direct absence of sleep (indeed, participants in the current study were slower to respond on the PVT and reported being less alert on the Stanford Sleepiness Scale after sleep deprivation than sleep). Focusing first on our assessment of overnight consolidation, generalised cognitive impairments arising from sleep deprivation could have impaired retrieval performance, creating the impression of a sleep-associated improvement in retention. While this is a reasonable concern in view of the sleep-memory effects observed at our delayed test (which followed immediately after the overnight interval), it does not explain why the retention advantage in the sleep condition was still present 48 h later (once sleep deprived individuals had had ample opportunity for recovery sleep). Moreover, we observed no significant relationship between the benefits of sleep (vs sleep deprivation) on memory retention and between-conditions differences in Stanford Sleepiness Scale scores or PVT response times, suggesting that our findings were not driven by the general cognitive impairments that accompany sleep deprivation. It is therefore reasonable to conclude that our data reflect a positive impact of sleep on memory consolidation. To what extent this memory benefit of sleep can be explained by an absence of wakeful interference (such as that experienced in the sleep deprivation condition) or an active sleep-dependent consolidation mechanism, however, cannot be inferred from our data.

Turning to our analysis of next-day learning, although the assessment phase also took place 48 h after encoding, the initial learning phase occurred immediately after sleep or sleep deprivation. We therefore cannot rule out the possibility that the apparent improvement in encoding performance after sleep was influenced by generalised cognitive impairments following sleep deprivation. Importantly, however, we think that our EEG data provide reasonable evidence that an absence of sleep does in itself disrupt new learning. Specifically, if our effects were driven by non-specific cognitive deficits following sleep deprivation, one would expect to have observed only generalised differences in EEG activity between the sleep and sleep deprivation conditions (i.e. only a main effect of condition, across all encoding trials). By contrast, a significant interaction showed that beta desynchronization was amplified in the sleep (vs sleep deprivation) condition, specifically on trials for which adjective-image pairings were subsequently remembered. This impact of sleep on beta desynchronization during successful learning was not predicted by between-condition differences in Stanford Sleepiness Scale scores or PVT response times, and no between-condition difference in beta power emerged for pairings that were subsequently forgotten (see Figures 4b and 4c). Because beta desynchronization is an established neural marker of semantic processing during successful learning (Griffiths et al. 2016; Hanslmayr et al. 2014; Hanslmayr et al. 2011), these findings may suggest that the neural mechanisms of encoding are indeed disrupted by an absence of sleep. Further support for this view is available below, where we outline how the brain may engage in compensatory learning strategies when semantic processing pathways are compromised by sleep deprivation (see *Sleep loss disrupts effective learning*).

Because our retention index was based on tests for the same items at the immediate, delayed and follow-up sessions, it is possible that our data were influenced by retrieval practice effects (i.e. memories that undergo retrieval practice are typically better remembered than those that do not, Carpenter et al. 2008; Roediger and Karpicke 2006). That is, the retention advantage observed after sleep (vs sleep deprivation) at the delayed test might have been maintained at the follow-up test as a result of retrieval practice. However, given that memories strengthened through retrieval gain little benefit from sleep-associated consolidation (Antony et al. 2017; Antony and Paller 2018; Bäuml et al. 2014), then, under a retrieval practice hypothesis, the immediate test should have nullified any later impact of sleep on retention. While it may still be argued that a between-condition difference in retention at the delayed test was driven by non-specific impairments following sleep deprivation, this would not explain why the memory advantage in the sleep condition was still present 48 h later (once recovery sleep had taken place). We therefore think that retrieval practice effects cannot provide a reasonable explanation of our findings.

Given that recovery sleep after sleep deprivation is characterised by a homeostatic increase in SWS (Borbély 1982; Borbély et al. 2016), one might have expected the overnight consolidation of newly learned adjective-image pairings to be amplified in the sleep deprivation (vs sleep) condition, potentially tempering the initial impact of sleep loss on encoding. Although we did not record sleep EEG during the time that participants were away from the laboratory (and thus have no insight into homeostatic increases in SWS after sleep deprivation), we did monitor sleep behaviour with wristwatch actigraphy. Participants slept for longer during the first night after learning in the sleep deprivation (vs sleep) condition, but this between-condition difference in sleep duration was not significantly correlated with the magnitude of sleep’s benefit for learning. This suggests that longer recovery sleep in the sleep deprivation condition did not meaningfully influence the impact of sleep loss on new learning.

It is worth noting, though, that an enhanced consolidation of newly learned adjective-image pairings in the sleep deprivation (vs sleep) condition (due to longer or deeper recovery sleep) could have obscured a relationship between sleep-associated memory retention and next-day learning in our multiple regression analysis. However, the same null effects were observed when our analysis was restricted to data from the sleep condition alone, rather than subtractions between the sleep and sleep deprivation conditions (as was done in our primary analysis). Hence, no relationship between overnight consolidation and next-day learning was observed when the influence of sleep deprivation (and the putative enhancement of sleep-associated consolidation during recovery sleep) was removed from our data.

### No link between sleep-associated consolidation and next-day learning

If memory consolidation during SWS supports a shift in the memory retrieval network from hippocampus to neocortex, then sleep-associated consolidation of hippocampus-dependent memories should predict next-day learning of new, hippocampally-mediated associations, and SWA should facilitate this relationship. However, we observed no such effects in our data, suggesting that new learning in hippocampus may not be contingent on hippocampal memory processing during the preceding night of sleep.

An alternative interpretation of these null effects is that our experimental paradigm could not provide an adequate test of our hypothesis. Although we reasoned that the use of two conceptually different hippocampus-dependent tasks would prevent our findings from being influenced by retroactive or proactive interference, qualitative differences between these tasks might have negated our ability to detect a relationship between sleep-associated memory consolidation and next-day learning. This is nevertheless a speculative suggestion that can be addressed in future research (e.g. by using a paired-associates task to assess both overnight memory retention and subsequent encoding).

Although our study was motivated by the assumptions of the Active Systems framework (Born and Wilhelm 2012; Klinzing et al. 2019; Rasch and Born 2013; Walker 2009), it is important to also consider our findings in the context of homeostatic synaptic downscaling, which is regarded as another fundamental mechanism through which sleep supports learning and memory (Tononi and Cirelli 2014; 2016). From this perspective, sleep is the price the brain pays for waking plasticity, in order to avoid an accumulation of synaptic upscaling. Because synaptic renormalization should mainly occur during sleep (when neural circuits can undergo a broad and systematic synaptic downscaling), a night of sleep deprivation would prevent the restoration of cellular homeostasis and impair next-day learning. A number of theoretical accounts of sleep-associated memory processing have made progress in reconciling the key tenets of the Active Systems and Synaptic Homeostasis frameworks, suggesting that these processes work in concert to support global plasticity and local downscaling, respectively, and in doing so prepare the hippocampus for future encoding (Genzel et al. 2014; Klinzing et al. 2019; Lewis and Durrant 2011). Interestingly, whereas global memory replay and consolidation have been linked to slow (< 1 Hz) oscillations, downscaling and forgetting are associated with delta waves (1-4 Hz) in local networks (Genzel et al. 2014; Kim et al. 2019). How interactions between global slow oscillations and local delta waves regulate overnight memory processing is therefore pertinent to further understanding of the relationship between sleep-associated consolidation and next-day learning.

### Sleep loss disrupts effective learning

Successful learning is associated with left-lateralised beta desynchronization ∼0.5-1.5s after stimulus onset (Griffiths et al. 2016; Griffiths et al. 2021; Hanslmayr et al. 2014; Hanslmayr et al. 2009; Hanslmayr et al. 2012; Hanslmayr et al. 2011). Consistent with these prior studies, we observed a decrease in beta power ∼0.3-2 s after stimulus onset during the encoding of subsequently remembered (vs forgotten) associations, and this was most pronounced in the left hemisphere. Beta desynchronization is thought to reflect semantic processing during successful memory formation (Fellner et al. 2013; Hanslmayr et al. 2011); as beta power decreases, the depth of semantic processing increases (Hanslmayr et al. 2009). More broadly, neocortical alpha/beta oscillations have been linked to the processing of incoming information during episodic encoding (Griffiths et al. 2021). For our learning task, participants were instructed to form vivid mental images or stories that linked the adjective and image of each pairing. The observed downregulation of beta power during successful learning might thus reflect an engagement of information processing operations, possibly involving semantic representations, allowing these novel associations to be bound together into one coherent episode and committed to memory.

Importantly, however, the change in beta power that accompanied successful learning differed according to whether participants had slept or remained awake across the overnight interval. Whereas encoding of subsequently remembered (vs forgotten) adjective-image pairings was associated with beta desynchronization after sleep, no significant difference in beta power emerged from the same contrast after sleep deprivation. Hence, a protracted lack of sleep appeared to disrupt semantic processing operations when participants were successfully forming new memories. This interpretation is in line with previous behavioural findings where sleep deprived individuals have had difficulty in encoding semantically incongruent stimulus pairs (Alberca-Reina et al. 2014). The sleep deprived brain might thus rely on alternative processing routes when committing new information to memory. Indeed, prior studies have shown that sleep deprivation leads to compensatory neural responses during learning (Chee and Choo 2004; Drummond et al. 2004) and recognition (Sterpenich et al. 2007).

What might be the nature of this alternative route to learning after sleep deprivation? It is interesting to note that we observed an upregulation of beta activity during successful (vs unsuccessful) learning in the sleep deprivation condition (although this difference did not survive a Bonferroni correction for multiple comparisons). Increases in beta power have been linked to working memory and active rehearsal (Deiber et al. 2007; Hwang et al. 2005; Onton et al. 2005; Tallon-Baudry et al. 2001), suggesting that sleep-deprived individuals may engage in more surface-based rehearsal strategies due to semantic processing pathways being compromised by an absence of sleep.

It is important to note that the foregoing findings on beta desynchronization arose from an exploratory analysis that was not pre-registered, and should therefore be treated with caution until such time that they are replicated in confirmatory research.

### Conclusions

We investigated whether memory consolidation in sleep predicts next-day learning and whether SWA contributes to this relationship. Furthermore, we investigated how the neural correlates of successful learning are affected by sleep deprivation. Although sleep improved both memory retention and next-day learning, we found no evidence of a relationship between these measures, nor with SWA. Whereas beta desynchronization – an established marker of semantic processing during successful learning – was present during the encoding of subsequently remembered (vs forgotten) associations after sleep, no such difference in beta power was observed after sleep deprivation. An extended lack of sleep might therefore disrupt our ability to draw upon semantic knowledge when encoding novel associations, necessitating the use of more surface-based and ultimately suboptimal routes to learning.

## Funding

This work was supported by a Medical Research Council Career Development Award (MR/P020208/1) to S.A.C and a University of York Department of Psychology Doctoral Studentship to A.áV.G.

## Acknowledgments

We are grateful to Marcus O. Harrington for help with data collection and Jennifer E. Ashton for help with the experimental materials. We also thank members of the Sleep, Language and Memory Group at the University of York for fruitful discussions of the data. Study data and analysis scripts can be retrieved via the following link: osf.io/cy2s9.

## Notes

https://osf.io/pjwrn/

### Competing Interest Statement

The authors have declared no competing interest.

### Summary of Updates

This version of the manuscript has been revised following peer review.

