## Supplementary Material for "Sleep loss disrupts the neural signature of successful learning"

**Psychomotor Vigilance Task**

A 2 (Condition: Sleep/Sleep Deprivation) x 3 (Session: Immediate/Delayed/Follow-up) repeated measures ANOVA with Greenhouse-Geisser correction showed that participants were significantly slower at responding in the sleep deprivation condition (F(1,28) = 20.71, p < .001 , ƞ_p_^2^ = 0.43) and that response times varied significantly across sessions (F(1.11,30.95) = 30.10, p < .001, ƞ_p_^2^ = 0.52). There was also a Condition * Session interaction (F(1.15,32.12) = 28.84, p < .001, ƞ_p_^2^ = 0.51), which was driven by slower responses in the morning after sleep deprivation as compared to sleep (Delayed Session: p < .001, Bonferroni corrected). Please note that one participant was removed from this analysis because their data from the follow-up session of the sleep deprivation condition was missing.

**Table 1.** Mean (± SEM) response times (ms) on the Psychomotor Vigilance Task at each session per condition. Note that one datapoint was missing for the follow-up session in the sleep deprivation condition.

| **Condition** | **Immediate Session** | **Delayed Session** | **Follow-up Session** |
| --- | --- | --- | --- |
| **Sleep** | 283.93 (± 5.05) | 289.15 (± 4.34) | 283.69 (± 6.11) |
| **Sleep Deprivation** | 278.03 (± 4.92) | 399.00 (± 17.63) | 287.46 (± 5.06) |

**Stanford Sleepiness Scale**

A 2 (Condition: Sleep/Sleep Deprivation) x 3 (Session: Immediate/Delayed/Follow-up) repeated measures ANOVA showed that participants rated themselves as feeling less alert in the sleep deprivation condition (F(1,29) = 72.95, p < .001, ƞ_p_^2^ = 0.72) and that their ratings varied significantly across sessions (F(2,58) = 85.95, p < .001, ƞ_p_^2^ = 0.75). There was also a Condition * Session interaction (F(2,58) = 82.34, p < .001, ƞ_p_^2^ = 0.74), which was driven by higher sleepiness ratings in the morning after sleep deprivation as compared to sleep (Delayed Session: p < .001, Bonferroni corrected).

**Table 2.** Mean (± SEM) ratings on the Stanford Sleepiness Scale at each session per condition.

| **Condition** | **Immediate Session** | **Delayed Session** | **Follow-up Session** |
| --- | --- | --- | --- |
| **Sleep** | 2.80 (± 0.15) | 2.27 (± 0.16) | 2.17 (± 0.11) |
| **Sleep Deprivation** | 2.63 (± 0.12) | 5.37 (± 0.15) | 2.20 (± 0.12) |
